# A selective adenylyl cyclase 1 inhibitor relieves pain without causing tolerance

**DOI:** 10.1101/2021.06.23.448127

**Authors:** Gianna Giacoletti, Tatum Price, Lucas V.B Hoelz, Abdulwhab Shremo Msdi, Katerina Vazquez-Falto, Tácio V. Amorim Fernandes, Vinícius Santos de Pontes, Hongbing Wang, Nubia Boechat, Adwoa Nornoo, Tarsis F. Brust

**Affiliations:** Department of Pharmaceutical Sciences, Lloyd L. Gregory School of Pharmacy, Palm Beach Atlantic University, West Palm Beach, FL, USA 33416; Drug Synthesis Laboratory, Drug Technology Institute, Farmanguinhos - FIOCRUZ, Oswaldo Cruz Foundation, Manguinhos, RJ, Brazil 21040900; Instituto Nacional de Metrologia, Qualidade e Tecnologia – INMETRO, Rio de Janeiro, RJ, Brazil 25250020; Department of Physiology, Michigan State University, East Lansing, MI, USA 48824

**Keywords:** Adenylyl cyclase, pain, analgesia, AC1, tolerance

## Abstract

Adenylyl cyclases (ACs) catalyze the production of the second messenger cyclic adenosine monophosphate from adenosine triphosphate. Among the ten different AC isoforms, studies with knockout animals indicate that inhibition of AC1 can relieve pain and reduce behaviors linked to opioid dependence. We previously identified ST034307 as a selective inhibitor of AC1. The development of an AC1-selective inhibitor now provides the opportunity to further study the therapeutic potential of inhibiting this protein in pre-clinical animal models of pain and related adverse reactions. In the present study we have shown that ST034307 relieves pain in mouse models of formalin-induced inflammatory pain, acid-induced visceral pain, and acid-depressed nesting. In addition, ST034307 did not cause analgesic tolerance after chronic dosing. We also show that the compound is restricted to the periphery following subcutaneous injections and report the predicted molecular interaction between ST034307 and AC1. Our results indicate that AC1 inhibitors represent a promising new class of analgesic agents that treat pain and appear to produce less adverse effects than currently-used opioids.

## Introduction

Adenylyl cyclases (ACs) are the enzymes responsible for catalyzing the conversion of adenosine triphosphate (ATP) into cyclic adenosine monophosphate (cAMP).^1,2^ ACs integrate signaling from a large range of proteins and ions, including G protein-coupled receptors (GPCRs), protein kinases, and calcium, to name a few. There are ten different isoforms of ACs, nine of them are present in the cellular membrane and one is soluble. Each AC isoform has a specific expression pattern, which is related to a specific set of physiological functions.^1^ AC isoforms also display a unique set of regulatory properties that result in differences in how the isoforms are modulated by different types of G proteins, protein kinases, and ions.^1,2^

AC1 is part of the group of ACs that are activated by calcium through calmodulin.^3^ Additional regulatory properties of AC1 include inhibition by Gα_i/o_ and Gβ_γ_ subunits of G proteins and activation by Gα_s_ and the small molecule forskolin.^1,4^ AC1 has also been shown to undergo Gα_i/o_-coupled receptor-mediated superactivation.^4–6^ The expression pattern of AC1 is consistent with the physiological functions that have been associated with this AC isoform. AC1 transcripts are found in the dorsal root ganglion (DRG), spinal cord, and anterior cingulate cortex (ACC), and a role for this cyclase in pain and nociception has been suggested.^7–9^ In fact, AC1 knockout (KO) mice display a reduction in typical behaviors that are induced by inflammatory and neuropathic pain, compared to wild-type mice.^8,10,11^ These studies encouraged the pursuit and discovery of novel compounds that can selectively inhibit AC1 activity as potential novel pain-relieving therapeutics.^4,12,13^

AC1 transcripts are also found in the hippocampus, a brain region linked to learning and memory.^14^ Notably, AC8, another calcium/calmodulin-activated isoform, is also highly expressed in the hippocampus.^1,15^ Previous studies with single and double AC1/AC8 KO mice have indicated that some functions of AC1 and AC8 related to learning and memory are redundant.^14^ Specifically, AC1/AC8 double KO mice display impaired long-term memory in contextual learning and passive avoidance assays, whereas individual KO of each isoform separately results in wild-type-like behaviors.^14^ However, each isoform also appears to have specific functions. While less severe deficits are observed in AC1-KO mice compared to the AC1/AC8 double KO, the former still displays reduced long-term potentiation (LTP) in the hippocampus and impairments in certain recognition memory as well as spatial and avoidance learning tasks.^16,17^ Those studies highlight the importance of selectivity for AC1 inhibition versus AC8 for a novel compound to treat pain, but do not exclude the possibility of adverse effects that may result from selective AC1 inhibition in the hippocampus.

We have recently reported the discovery of ST034307, a small molecule inhibitor of AC1 that is selective for AC1 inhibition versus all other membranous AC isoforms, including AC8.^4^ Our previous study focused on the characterization of ST034307 at the molecular level, showing that the compound is a potent, highly selective, and direct AC1 inhibitor. Moreover, ST034307 was also analgesic in a mouse model of Complete Freund’s Adjuvant (CFA)-induced allodynia.^4^ The present study represents a pre-clinical study with ST034307 to determine the potential of this class of compounds as novel analgesic agents. We compared the compound with morphine in mouse models of pain-induced and pain-depressed behaviors and also showed that the compound is restricted to the periphery following subcutaneous injections. Further, we showed that ST034307 does not induce analgesic tolerance or cross-tolerance with morphine. Finally, we expanded our previous mechanistic findings by modeling how the interaction of ST034307 with AC1 happens.

## Materials and Methods

### Experimental design

The main goal of the present study was to determine the potency and efficacy of the AC1 selective inhibitor ST034307 in mouse models of pain and innate behavior. Male mice were used as research subjects and sample sizes for the different experiments were determined using power analyses from preliminary experiments following the guidelines of Palm Beach Atlantic University’s Institutional Animal Care and Use Committee (IACUC) to attempt to minimize the numbers of animals used. Instances where the number of animals per group vary in an experiment were the result of additional animals being required for proper blinding when a drug dose had to be added. All animals were randomized to treatments and experimenters performing behavioral measurements and injections were blinded to all compound treatments and doses.

### Materials

ST034307 (6-Chloro-2-(trichloromethyl)-4H-1-benzopyran-4-one) was purchased from Tocris Bioscience and morphine sulphate from Spectrum Laboratory Products. Acetic acid, lactic acid, Tween 80, and formaldehyde were from Sigma-Aldrich. Dimethyl sulfoxide (DMSO) and 0.9% sterile saline were from Fisher Scientific. ST034307 and morphine were prepared in a vehicle consisting of dimethyl sulfoxide (DMSO), Tween 80, and milli-Q water (1:1:8). Specifically, ST034307 was first dissolved in DMSO and sonicated in a 50°C water bath for 15 minutes. Next, Tween 80 was added, the solution was vortexed, and the sonication was repeated. Warm (37°C) milli-Q water was added and the solution was vortexed immediately before injections. Acetic acid, lactic acid, and formalin were diluted in 0.9% sterile saline.

### Animals

Male C57BL/6J mice were purchased from Charles Rivers Laboratories. AC1-KO mice were created as previously described and propagated using homozygous breeding using the C57BL/6J background.^17^ This particular strain is commonly used in studies related to analgesic agents^18–20^ and provides a way of comparing the activity of ST034307 with other compounds, given that morphine was used as a positive control. Mice were housed in groups (2-5 per cage) in cages covered with filter tops (micro barrier top from Allentown), in a temperature-controlled room under a 12-hour light/dark cycle. Animals had *ad libitum* access to water and food, as well as nesting material made from pulped virgin cotton fiber (nestlets from Lab Supply) for enrichment. Corn cob bedding (1/4”) was used for bedding. Mice between 2 and 5 months of age were used for experiments and were dosed subcutaneously with 10 μl/g of ST034307, morphine, or vehicle solutions. After each experiment, mice were humanely euthanized via cervical dislocation under isoflurane anesthesia (open drop method).

### Study approval

All experimental procedures involving mice adhered to the National Institutes of Health Animal Care guidelines and were approved by Palm Beach Atlantic University’s IACUC (West Palm Beach, FL).

### Formalin-induced paw licking

The formalin-induced paw licking assay was conducted similarly to previously described.^20^ Briefly, mice were acclimated to clear testing cylinders for 45 minutes. Next, mice were injected subcutaneously with compounds or vehicle solutions and returned to acrylic cylinders for 15 minutes. Mice were then injected into their right hind paw with 25 μl of 5% formalin using a 25 μl Hamilton syringe and a 30-gauge needle. Mice were immediately returned to the testing cylinders, and paw licking time was recorded in 5-minute intervals for 40 minutes. The experiment was divided into two different phases. The first represents the time spent licking between 0 and 10 minutes, the second represents the time spent licking between 16 and 40 minutes.

### Acid-induced writhing

For the acid-induced writhing assay, mice were acclimated to clear testing cylinders for 45 minutes. Next, mice were injected subcutaneously with compounds or vehicle solutions and returned to acrylic cylinders for 15 minutes. Mice were then injected intraperitoneally with 0.75% acetic acid (10 μl/g), returned to the testing cylinders, and the number of abdominal constrictions (stretching movements of the body as a whole, including the hind paws) was counted in 5-minute intervals for 30 minutes as previously described.^21^ For the tolerance assay, mice were injected subcutaneously with either 100 mg/kg morphine or 30 mg/kg ST034307 (solubility issues prevented the use of higher doses) once a day for four or eight days. At day four or day eight, acid-induced writhing assays were performed.

### Nesting

The mouse nesting assay was adapted from methods previously described.^22^ Mice were single housed and acclimated to their new home cage for three days. During the following three days, mice underwent one nesting session (as described below) per day to acclimate them to handling, the experimental procedure, and the testing room. The last acclimation session included a subcutaneous injection (for compound-inhibited nesting) or a subcutaneous injection and an intraperitoneal injection (for acid-depressed nesting) with 0.9% saline. On the day after the third acclimation session, mice were injected subcutaneously with compounds or vehicle and returned to their respective home cages for 10 minutes. Mice were transferred to a transfer cage (< 1 minute) and nestlets we placed in each of the 6 different zones of the home cage as previously described.^22^ Mice were either returned to their home cages (compound-inhibited nesting) or injected intraperitoneally (10 μl/g) with 1% lactic acid (acid-depressed nesting) and returned to their home cages for nesting periods. Nesting was scored as the number of zones cleared over time.

### Pharmacokinetic studies

The disposition of ST034307 was studied in male C57BL/6J mice following a single subcutaneous injection (10 mg/kg). Mice were humanely euthanized via decapitation under isoflurane anesthesia (open drop method). Subsequently, brain and blood samples were collected in triplicate at 5-, 25-, 45-, 60-, 120- and 240-minutes post-injection. Blood was centrifuged, plasma collected and stored at −80°C. The analyses of the samples were conducted in the Drug Metabolism and Pharmacokinetics Core at Scripps Research. Brain samples were homogenized with water to form a slurry. ST0304307 was extracted from plasma and brain slurry on solid-supported liquid-liquid extraction cartridges (HyperSep™, SLE, 1g/6ml, Thermo Scientific) and the resultant extract was assayed for ST034307 by tandem mass spectroscopy coupled to HPLC (SCIEX 6500). A plot of plasma ST0304307 concentration versus time was constructed and analyzed for non-compartmental pharmacokinetic parameters - half-life, volume of distribution and clearance (Phoenix, Pharsight, Certara Inc.).

### Molecular Docking

#### Construction of the AC1 model

The AC1 model was constructed through ab-initio and threading methods on I-Tasser server, considering as input the sequences Phe291-Pro478 and Leu859-1058, registered under the UniProtKB ID Q08828.^23,24^ The globular domain regions were identified using both Pfam and UniProtKB feature viewer, being selected for further refining.^25^ Local sequence alignments with NCBI’s BLAST+ were made between the Q08828 and those from Protein Data Bank (PDB) deposited structures to find experimentally solved structures with magnesium ions, ATP, and Gα_s_ on their respective sites.^26,27^ Thus, using molecular superpositions on VMD 1.9.3, the cofactors and ligands were extracted from the structure registered as 1CJK on PDB, while Gα_s_ was extracted from the structure registered as 6R3Q, and positioned into the AC1 model.^28^ MODELLER 9.25-1 was then used to run 100 cycles of structural optimizations with molecular dynamics, simulated annealing, and conjugated gradient.^29^ The generated structures were ranked by DOPE-Score and the best model was selected. To verify the structural quality of the best AC1 model built, the structure was submitted to the SAVES server, where two programs were selected, PROCHECK (Figs. S1, S2, and S3) and VERIFY 3D (Fig. S4), and to the Swiss-PROT server, using QMEANDisCo algorithm (Fig. S5).^30–32^

#### Preparation of the ST034307 structure

ST034307 was constructed and optimized with the HF/6-31G(d) level of theory using the SPARTAN’16 program (Wavefunction, Inc.).

#### Docking using GOLD 2020.3.0 (Genetic Optimization for Ligand Docking)

The molecular docking simulation using the GOLD program was carried out using automatic genetic algorithm parameters settings for the population size, selection-pressure, number of islands, number of operations, niche size, and operator weights (migration, mutation, and crossover).^33^ The search space was a 40 Å radius sphere from the 66.215, 105.567, and 81.040 (x, y, and z axes, respectively) coordinates. The scoring function used was ChemPLP, which is the default function for the GOLD program. Thus, the pose with the most positive score (the best interaction) was extracted for further analysis.

#### Docking using AutoDock Vina 1.1.2

The PDBQT-formatted files for the AC1 model and ST034307 structure were generated using AutoDock Tools (ADT) scripts.^34^ Using the AutoDock Vina program, the grid size was set to 65.172 Å × 77.050 Å × 73.559 Å for x, y, and z axes, respectively, and the grid center was chosen using 66.215 (x), 105.567 (y), and 81.040 (z) as coordinates. Each docking run used an exhaustiveness setting of 16 and an energy range of 3 kcal/mol. Consequently, the pose with the lowest energy was extracted for interaction analysis.

### Data and statistical analyses

All statistical analyses were carried out using GraphPad Prism 9 software (GraphPad Software Inc.). Data normalization and nonlinear regressions were carried out similarly to previously described.^19^ For normalizations (representing a rescaling of the Y axis for enhanced clarity), the maximal possible effect was set as 100% (zero for formalin-induced paw licking and acid-induced writhing, and five for acid-depressed nesting) and the response to vehicle’s average as 0%. For compound-inhibited nesting, the response from vehicle’s average was defined as 100% and zero to 0%. Normalized data was fitted to three-parameter nonlinear regressions with the top constrained to 100% and the bottom to 0% (except for the cases where ST034307 did not reach a full response, where no top constrain was set). All statistical analyses were performed using raw experimental data (without normalization). T tests with Welch’s correction were used for comparisons between genotypes, one-way ANOVAs for comparisons within groups, and two-way ANOVAs for time-course evaluations. All ANOVAs where F achieved a statistical level of significance (*p* < 0.05) were followed by Dunnett’s corrections.

## Results

### ST034307 relieves inflammatory pain, but not acute nociception in mice

We have previously shown that intrathecal administration of ST034307 relieves CFA-mediated allodynia in mice.^4^ Here, we used intraplantar formalin injections to the mice’s right hind paws and compared the potency of ST034307 with that of morphine (both administered subcutaneously) for diminishing acute nociception and relieving inflammatory pain. The time spent tending to (licking) the injected paw was recorded (Figs. 1a and 1b). As indicated by previous studies using AC1-KO mice,^11^ only morphine caused a significant reduction in acute nociception, with an ED_50_ value equal to 5.87 mg/kg [95% CI 0.44 to 8.96] (Fig. 1c - sum of measurements recorded between 0 and 10 minutes). No significant effect was observed with ST034307. In contrast, both compounds significantly reduced formalin-induced paw licking in the inflammatory pain phase of the model compared to vehicle and had ED_50_ values equal to 6.88 mg/kg [95% CI 0.85 to 14.05] and 1.67 mg/kg [95% CI 0.35 to 2.43] for ST034307 and morphine, respectively (Fig. 1d – sum of measurements recorded between 16 and 40 minutes).

**Fig. 1.**
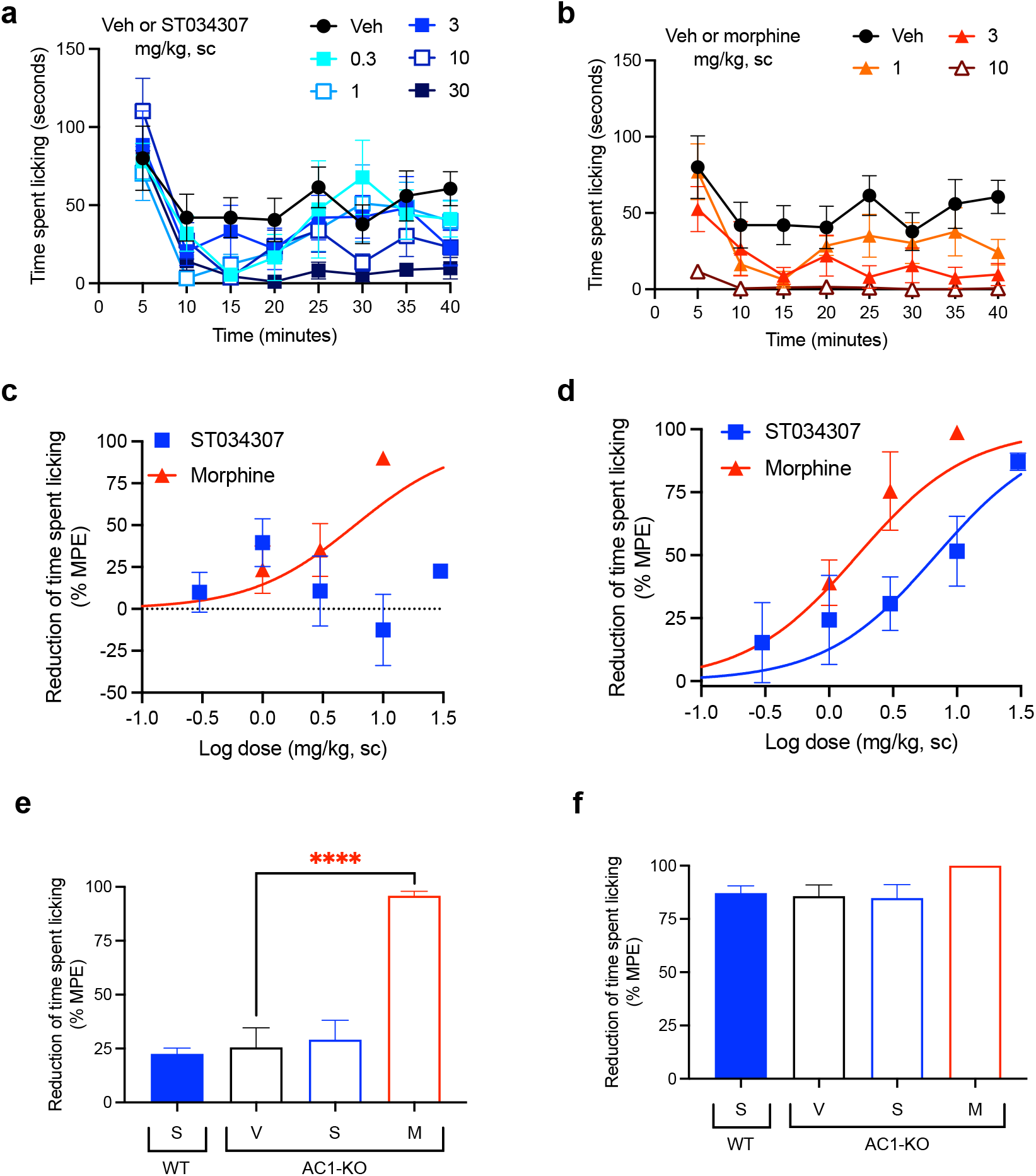
ST034307 relieves inflammatory pain in mice. (**a**) Different doses of ST034307 reduce paw licking behavior caused by an intraplantar injection with 5% formalin. (**b**) Different doses of morphine reduce paw licking behavior caused by an intraplantar injection with 5% formalin. (**c**) Dose-response curves of the sum of time spent licking the paw during the first 10 minutes of the graphs in A and B. Vehicle’s response was set as 0% and the maximal possible effect (0) to 100%. (**d**) Dose-response curves of the sum of time spent licking the paw during the period in between minute 16 and minute 40 of the graphs in A and B. Vehicle’s response was set as 0% and the maximal possible effect (0) to 100%. (**e**) Reduction of time spent licking the injected paw in wild-type (WT) and in AC1-KO mice treated with vehicle (V), 30 mg/kg ST034307 (S), or 10 mg/kg morphine (M) during the first 10 minutes of the experiment. (**f**) Reduction of time spent licking the injected paw in wild-type (WT) and in AC1-KO mice treated with vehicle (V), 30 mg/kg ST034307 (S), or 10 mg/kg morphine (M) during the first 10 minutes of the experiment. For E and F vehicle’s response in wild-type mice was set to 0% and zero to 100%. Data in all graphs represent the average ± S.E.M., N = 6-8. \*\*\*\**p* < 0.0001 in one-way ANOVA with Dunnett’s test.

Consistent with the results from wild-type mice, AC1-KO mice did not present a reduction of licking during the acute nociception phase of the experiment compared to wild-type mice (*p* = 0.2089 in unpaired t test – Fig. 1e). In addition, while morphine relieved acute nociception in AC1-KO mice (*p* < 0.0001 in one-way ANOVA), no effects were observed with ST034307 (Fig. 1e). In contrast, AC1-KO mice displayed a significant reduction of licking in the inflammatory phase of the model, compared to wild-type mice (*p* < 0.001 in unpaired t test – Fig. 1f). That reduction was similar to the effect 30 mg/kg ST034307 had in wild-type mice (Fig. 1f). No effects were observed from a dose of 30 mg/kg ST034307 in AC1-KO mice. Morphine (10 mg/kg) had a small effect in the inflammatory phase, but it was not significantly different from vehicle (*p* = 0.087 in one-way ANOVA – Fig. 1e).

### ST034307 relieves visceral pain and does not induce analgesic tolerance in mice

Visceral pain was induced by an intraperitoneal injection of 0.75% acetic acid. The number of abdominal stretches (writhing) the mice performed over a period of 30 minutes was recorded (Figs. 2a and 2b). Both ST034307 and morphine significantly reduced acid-induced writhing in this model with ED_50_ values equal to 0.92 mg/kg [95% CI 0.15 to 4.41] and 0.89 mg/kg [95% CI 0.40 to 1.52], respectively (Fig. 2c). However, ST034307 did not reach full efficacy at doses up to 30 mg/kg. Similarly, AC1-KO mice also only showed a partial reduction of acid-induced writhing in this model, compared to wild-type mice (*p* < 0.001 in unpaired t test, Fig. 2d). This response was not enhanced by 10 mg/kg ST034307, but 3 mg/kg morphine caused a significant reduction of acid-induced writhing in AC1-KO mice (*p* < 0.01 in one-way ANOVA – Fig. 2d).

**Fig. 2.**
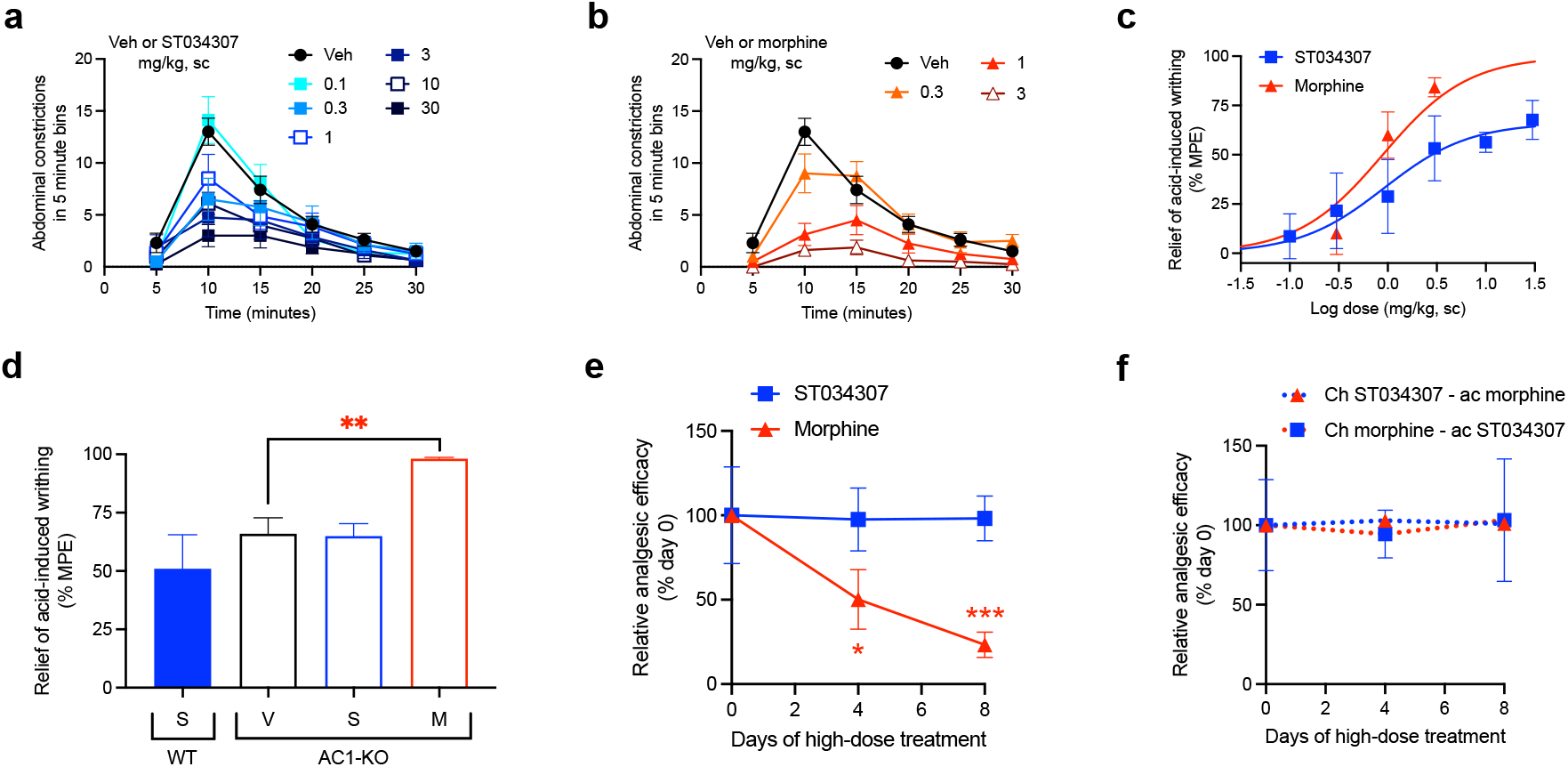
ST034307 relieves visceral pain in mice. (**a**) Different doses of ST034307 reduce the number of abdominal constrictions caused by an intraperitoneal injection with 0.75% acetic acid; N = 8-10. (**b**) Different doses of morphine reduce the number of abdominal constrictions caused by an intraperitoneal injection with 0.75% acetic acid; N=8-10. (**c**) Dose-response curves of the total number of abdominal constrictions from the graphs in A and B. Vehicle’s response was set as 0% and the maximal possible effect (0) to 100%. (**d**) Reduction of the total number of abdominal constrictions in wild-type (WT) and in AC1-KO mice treated with vehicle (V), 30 mg/kg ST034307 (S), or 10 mg/kg morphine (M). Vehicle’s response in wild-type mice was set as 0% and zero to 100%; N = 5. (**e**) Mice were injected daily with 30 mg/kg ST034307 or 100 mg/kg morphine and on day four or eight acid-induced writhing assays were performed. Mice that chronically received morphine displayed a decrease in efficacy of 3 mg/kg morphine on days four or eight, compared to a group of mice that received vehicle plus 3 mg/kg morphine on day zero. Mice that chronically received ST034307 did not display any changes in efficacy of 10 mg/kg ST034307; N = 5. (**f**) Mice were injected daily with 30 mg/kg ST034307 or 100 mg/kg morphine and on day four or eight acid-induced writhing assays were performed. Mice that chronically received morphine did not display any changes in efficacy with 10 mg/kg ST034307. Mice that chronically received ST034307 did not display any changes in efficacy with 3 mg/kg morphine; N = 5. Data in all graphs represent the average ± S.E.M. \**p* < 0.05, \*\**p* < 0.01, *** *p* < 0.001 in one-way ANOVA with Dunnett’s test.

Mice treated chronically with morphine display analgesic tolerance. Tolerance is expressed through the gradual loss in efficacy of a compound’s dose over time.^35^ After four days of daily subcutaneous injections with 100 mg/kg morphine, the efficacy of a 3 mg/kg dose of morphine decreased by nearly half (Fig. 2e). At day eight, morphine’s efficacy was nearly 20% of its initial response (Fig. 2e). In contrast, daily subcutaneous injections with 30 mg/kg ST034307 (highest dose we were able to inject chronically due to solubility) caused no decrease in the analgesic efficacy of a 10 mg/kg ST034307 dose at day four or day eight (Fig. 2e). Notably, no cross-tolerance was developed between the two compounds (Fig. 2f). Mice treated daily with 100 mg/kg morphine were still fully responsive to 10 mg/kg ST034307 at days four and eight; and mice treated daily with 30 mg/kg ST034307 were also fully responsive to 3 mg/kg morphine at days four and eight (Fig. 2f).

### ST034307 promotes analgesia in the absence of disruptions in the mouse nesting model

Nesting is an innate mouse behavior that can be disrupted by a number of different stimuli.^22^ Drugs, stress, and pain can all impede normal nesting behavior, making the model appropriate for detecting possible adverse reactions.^36^ In the experiment, nesting material (nestlets) was placed in six different zones of a mouse’s cage. As the mouse makes its nest, it gathers all nestlets in a single zone.^22^ We measured the numbers of zones cleared over time. ST034307 did not disrupt nesting behaviors at doses up to 30 mg/kg compared to vehicle (Fig. 3a). Morphine, on the other hand, caused a robust reduction of nesting behavior at 3 mg/kg (Fig. 3b – two-way ANOVA, *p* < 0.001 and *p* < 0.01 at 30 and 60 minutes, respectively). Morphine’s disruption of nesting behavior at the last time-point of the experiment resulted in an ED_50_ equal to 3.04 mg/kg [95% CI 1.16 to 11.32] (Fig. 3c).

**Fig. 3.**
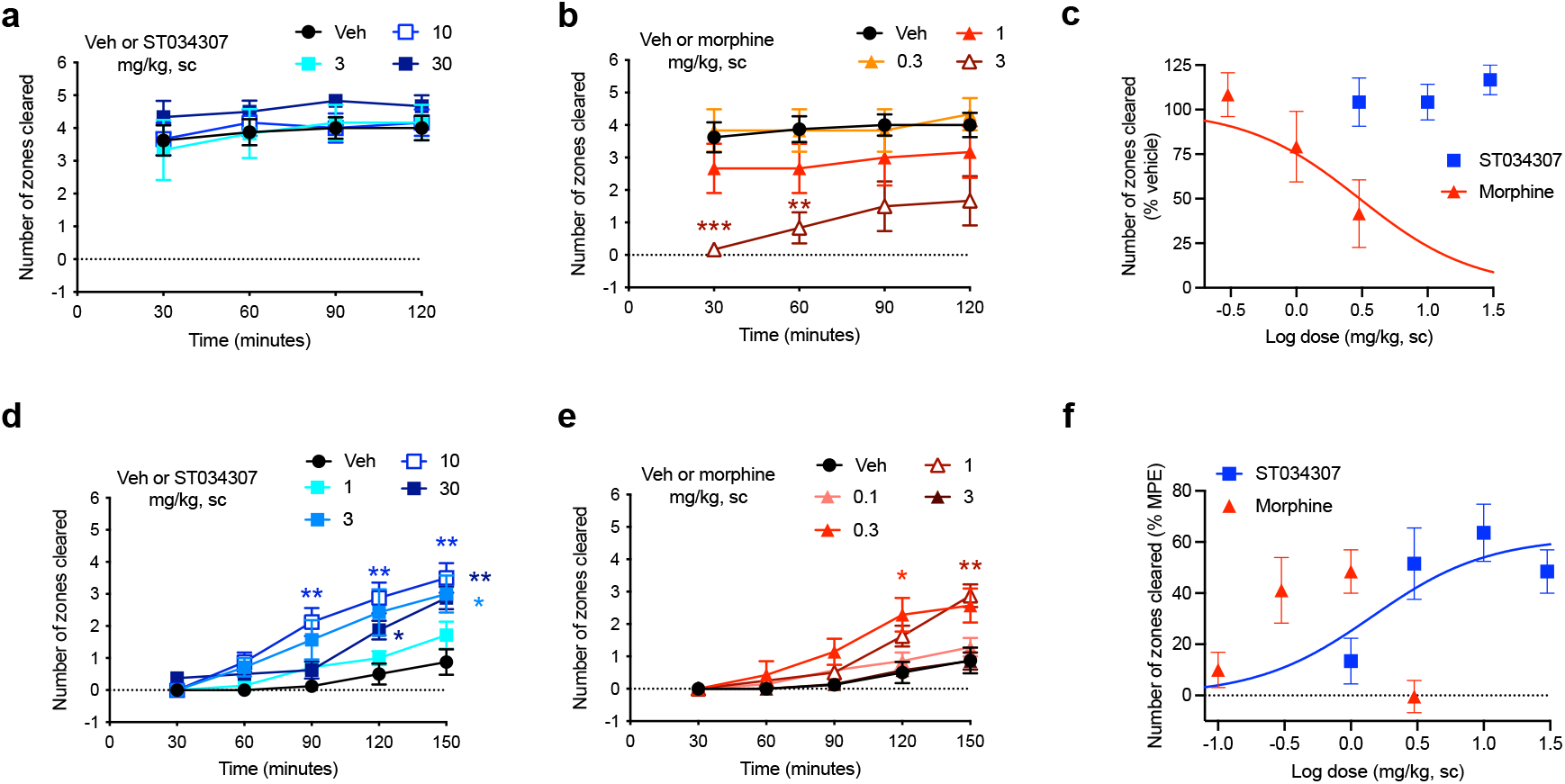
ST034307 rescues acid-depressed mouse nesting behavior. (**a**) ST034307 did not reduce mouse nesting behavior at doses up to 30 mg/kg. (**b**) At a dose of 3 mg/kg, morphine significantly reduced mouse nesting behavior at 30 and 60 minutes after nesting measurements were started. (**c**) Dose-response curves of the last experimental timepoints the graphs in A and B. Vehicle’s response was set as 100% and the maximal possible effect (0) to 0%. (**d**) and (**e**) 3 mg/kg, 10 mg/kg, and 30 mg/kg ST034307 and 1 mg/kg morphine significantly rescued mouse nesting behavior that was reduced by an intraperitoneal injection of 1% lactic acid. (**f**) Dose-response curves of the last experimental timepoints from the graphs in D and E. Vehicle’s response was set as 0% and the maximal possible effect (5) to 100%. Data in all graphs represent the average ± S.E.M., N = 6-8. \**p* < 0.05, \*\**p* < 0.01, \*\*\**p* < 0.001 in two-way ANOVA with Dunnett’s test.

As pain can also disrupt nesting behavior, we next tested whether ST034307 could recover nesting in mice that were treated with 1% lactic acid intraperitoneally. Lactic acid treatment caused a profound reduction in nesting behavior (Figs. 3d and 3e). Mice that were treated with 3, 10, or 30 mg/kg ST034307 displayed a significant increase in nesting behavior compared to vehicle-treated animals (Fig. 3d). For morphine, 0.3 and 1 mg/kg caused a significant recovery of nesting behavior during the assay, with no significant effects from 0.1 or 3 mg/kg (Fig. 3e). ST034307’s recovery of nesting behavior at the last time-point of the experiment resulted in an ED_50_ equal to 1.45 mg/kg [95% CI 0.22 to 4.93] (Fig. 3f). As the 3 mg/kg dose of morphine depressed mouse nesting, an ED_50_ value was not calculated for the compound (Fig. 3e). The ED_50_ value and partial response of ST034307 in this experiment are consistent with what was observed in the acid-induced writhing assay (Fig. 2c).

### ST034307 is restricted to the periphery

Given the positive results from the nesting experiments, we decided to determine the concentrations of ST0340307 in plasma and brain of mice at different timepoints following a subcutaneous injection with a dose of 10 mg/kg (Fig. 4). A plasma concentration of 0.44 (± 0.08) μM was observed immediately following the injection at 5 minutes. A sharp peak was present 60 minutes after the injection at 1.82 (± 0.39) μM and after 90 minutes, the plasma concentration dropped back to levels similar to the levels before the peak (0.33 μM ± 0.09). The half-life of ST0304307 was determined to be approximately 161 (± 88) minutes and the compound was rapidly cleared (CL/F) from the body at a rate of 305.04 (± 22.63) ml/min. ST0304307 may be highly tissue bound as its volume of distribution (V/F) of 1619 (± 790) ml is much greater than the total body water volume (14.5 ml) of a 24 g (average weight) mouse.^37^ This type of distribution may also indicate extensive red blood cell uptake. To our surprise, none of the timepoints measured resulted in detectable levels of ST034307 in the brains of those mice. These data indicate that the analgesic properties of ST034307 observed are due to its actions in the periphery.

**Fig. 4.**
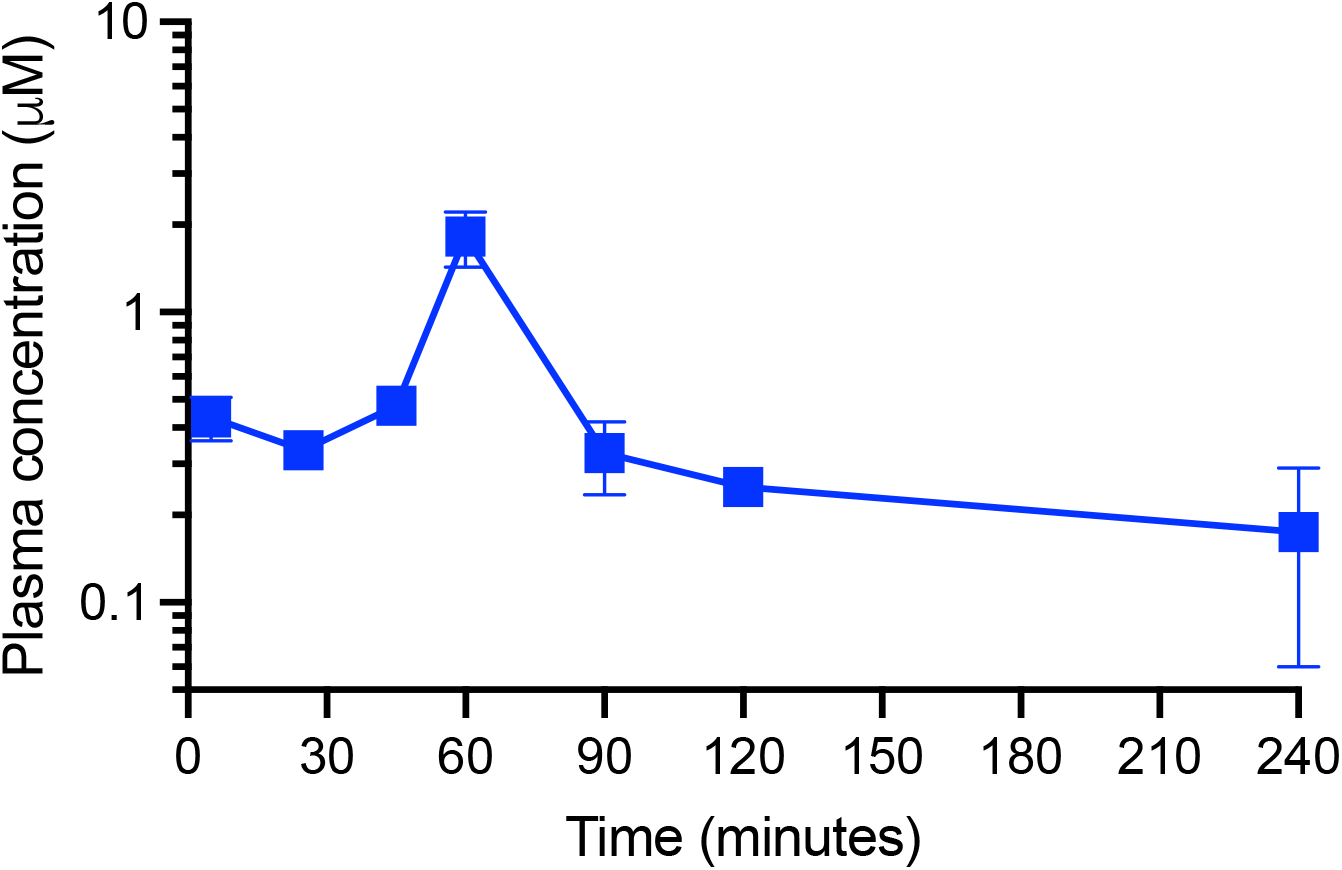
Plasma ST0304307 concentration versus time profile after a single subcutaneous injection in mice. Mice were injected with a dose of 10 mg/kg. Data in the graph represent the average ± S.E.M., N = 2-3.

### ST034307 interacts with the interface of C1a and C2a domains of AC1

In order to determine the binding interaction of ST034307 with AC1, we constructed a molecular model of AC1. The results of PROCHECK’s Ramachandran regions (Fig. S1), main-chain (Fig. S2), and side-chain parameters (Fig. S3), as well as VERIFY 3D (Fig. S4), and QMEANDisCo (Fig. S5) analyses indicated that the AC1 model was structurally valid to further computational studies. Thus, to predict the binding mode of ST034307 to AC1, we carried out molecular docking simulations using two programs, GOLD 2020.3.0 and Autodock Vina 1.1.2.^33,34^ Although these programs present differences concerning their search algorithm and scoring function, the best-predicted poses resulting from the different programs showed similar binding modes (RMSD = 2.35Å) into the AC1 model (Fig. 5a). The binding site was located into a cavity adjacent to the ATP binding pocket and between domains C1a and C2a, at the catalytic site interface. The best predicted pose for ST034307 presents a chemPLP score of 49.36 a.u, using the GOLD software, showing a hydrogen bond with the amine group from the side chain of Lys920 (C2a), and steric interactions with Phe306, Leu350, Cys353, Tyr355, Asp417, Val418, Trp419, Val423, Asn427, and Glu430 from C1a and with Lys920 and Ile922 from C2a (Figs. 5b and S6a). Using Autodock Vina, the best-predicted pose for ST034307 presents an interaction energy value of −6.9 kcal/mol, showing only steric interactions with Phe306, Leu350, Cys353, Tyr355, Asp417, Val418, Trp419, Ser420, Val423, Thr224, and Asn427 from C1a and with Lys920 and Ile922 from C2a (Figs. 5c and S6b).

**Fig. 5.**
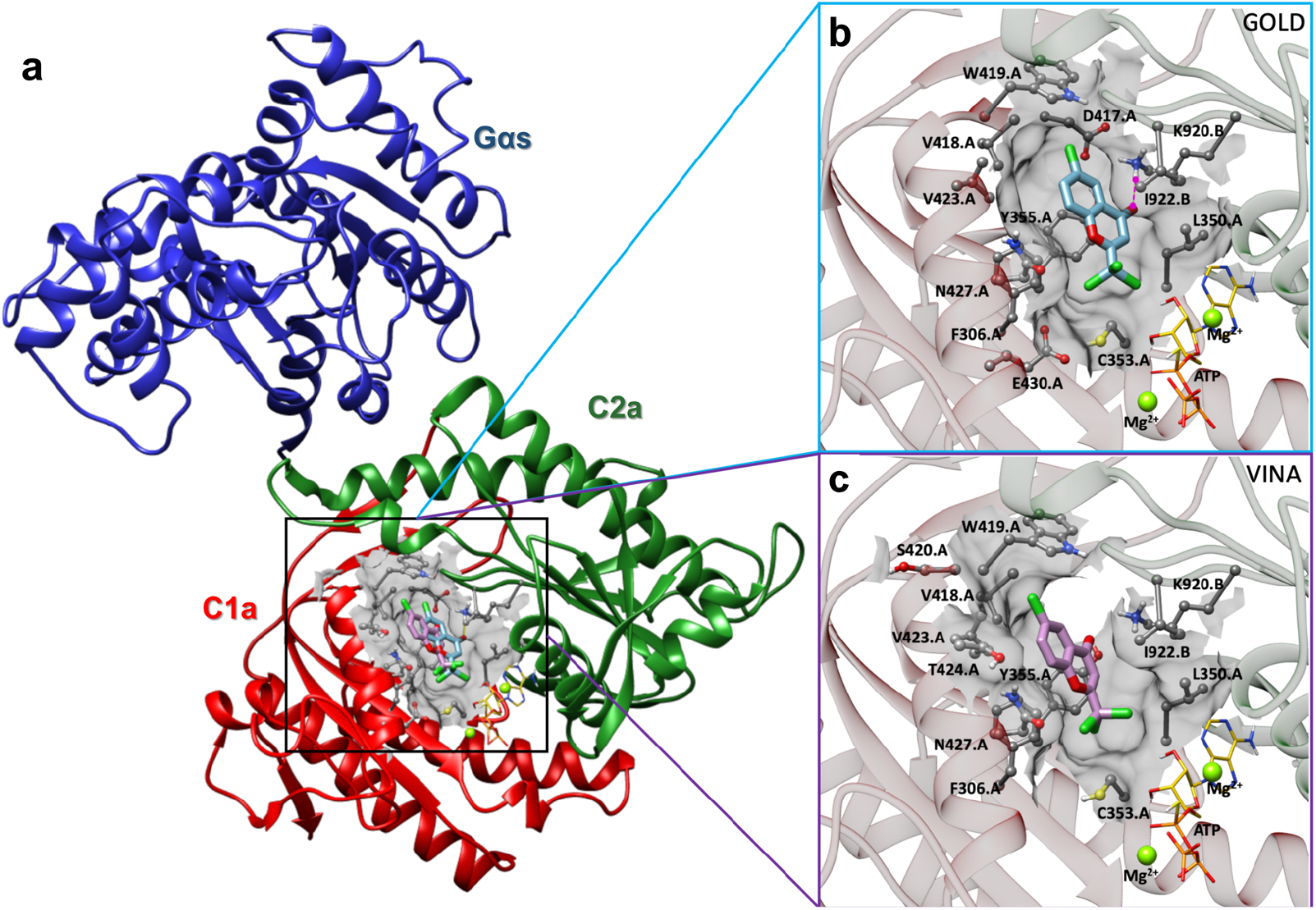
Prediction of the interaction between AC1 and ST034307. **(a)** Cartoon representation of the AC1 model, showing its catalytic domain (C1a, in red, and C2a, in green) complexed to Gα_s_ (in blue), ST034307 (in cyan or purple), ATP (in yellow), and two magnesium ions (Mg^2+^, in green). Predicted poses of ST034307, using Gold **(b)** and Autodock Vina **(c)** programs, presenting hydrogen-bond (interrupted purple line) and steric interactions. The AC1 residue structures are shown as ball and stick models, the ST034307 inhibitor and ATP as stick models, and Mg^2+^ ions as sphere models using UCSF Chimera program.^47^ All the structures are colored by atom: the nitrogen atoms are shown in blue, the oxygen atoms in red, the chlorine atoms in green, the hydrogen atoms in white, and the carbon chain in gray, cyan, or purple. Non-polar hydrogens have been omitted for clarity.

## Discussion and Conclusions

Previous studies using AC1-KO mice have indicated that inhibition of AC1 could be a new strategy to treat pain and opioid dependence.^8,10,11,38,39^ Inspired by those studies, we discovered and characterized ST034307.^4^ The compound displayed remarkable selectivity for inhibition of AC1 vs. all other membrane-bound AC isoforms. And while our previous manuscript focused on the molecular characterization of ST034307, we also showed that the compound relieves pain in a mouse model of CFA-induced allodynia.^4^ Here, those findings were expanded in multiple different ways.

First, we focused on the activity of the compound in two different models of pain-induced behaviors. In the first, intraplantar injections with formalin to the hind paws of the mice induce a paw licking behavior that is reflective of pain.^40^ The experiment is divided into two distinct phases. The first phase, which includes the first 10 minutes, represents chemical nociception due to the action of formalin on primary afferent nerve fibers.^41^ ST034307 had no effects on that phase of the experiment (Fig. 1c). This is consistent with our results with AC1-KO mice (Fig. 1e) and with a previous study that showed that AC1-KO mice do not have increased thresholds to thermal, mechanical, or chemical acute nociception compared to wild-type mice.^11^ Morphine, in contrast, reduced chemical nociception in both wild-type and AC1-KO mice. The formalin-induced paw licking behavior between minutes 16 and 40 is believed to be caused by the development of an inflammatory reaction that induces nerve sensitization.^36,40,42^ As expected, ST034307 caused a reduction in licking behavior during that phase. A reduction of formalin-induced paw licking during that phase was also observed in AC1-KO mice compared to wild-type animals. These data are consistent with previous work showing that AC1-KO mice have an increased threshold to inflammatory pain and indicate a possible use of selective AC1 inhibitors to treat this type of pain.^11^

Next, we showed that ST034307 decreases the number of abdominal constrictions (writhing) in mice injected intraperitoneally with acetic acid. Intraperitoneal injections with irritant agents cause peritovisceral pain and previous studies suggest that all analgesics can reduce abdominal cramps in this model.^36,43^ In contrast to morphine, ST034307 did not result in the maximal possible effect in this experiment, an outcome that was mimicked by AC1-KO mice. This partial response allowed us to further confirm that the effect of ST034307 in this model was through AC1 inhibition, as morphine, but not ST034307, further reduced the number of acid-induced abdominal constrictions in AC1-KO mice (Fig. 2d).

The use of analgesic agents often requires chronic dosing, which may last days, months, or even years depending on the patient’s pain condition. Unfortunately, chronic analgesic dosing may lead to analgesic tolerance.^44^ Opioid tolerance is well documented in humans and rodents, and results in a loss of analgesic efficacy over time.^19,35,44^ At the molecular level, it has been proposed that opioid tolerance is caused by agonist-induced recruitment of βarrestins to the mu opioid receptor (MOR). βarrestins induce receptor internalization (removal from the membrane) and, therefore, reduce the pool of available receptors for opioid action.^35^ As ST034307 acts as an inhibitor of AC1, the mechanisms commonly linked to tolerance (receptor downregulation) should not be present. Consistently, we did not observe any tolerance to a high daily dose of ST034307 for up to eight days in the mouse acid-induced writhing assay. This is in contrast to morphine, which displayed a marked reduction of analgesic efficacy, consistent with analgesic tolerance. As the two compounds act through different mechanisms (though the MOR inhibits AC1),^4^ there was no observable development of cross-tolerance.

Paw licking and abdominal constrictions are examples of pain-stimulated behaviors. While useful in pain studies, a reduction of these behaviors may not necessarily indicate pain relief. Compounds that induce paralysis, sedation, or stimulate a competing behavior, for instance, can still cause a marked reduction of behavior in those experiments, but are not necessarily relieving pain.^22,36^ Therefore, we have employed the pain-depressed behavior of nesting as another method to determine the analgesic efficacy of ST034307. Different types of stimuli (such as pain, stress, and sedation) can cause disruptions of mouse innate behaviors. Therefore, in order for a compound to display pain relief in this model, it may not present disruptive properties, as if it does, nesting behavior will be further reduced (see the 3 mg/kg dose of morphine in Figs. 3c and 3e).^22^ ST034307 did not disrupt nesting behavior at doses up to 30 mg/kg, indicating good tolerability in this model. Furthermore, all doses that were effective at relieving pain in the previous models, also significantly recovered nesting behavior that was reduced by an intraperitoneal injection of lactic acid (Fig. 3d). According to Negus (2019), the combination of the results from our nesting experiments and our pain-stimulated behavior experiments makes ST034307 (and possibly other AC1-selective inhibitors) a “high-priority” analgesic compound for “further testing”.^36^

While the nesting experiments provide a measure of safety, studies describing the full spectrum of possible adverse reactions that result from AC1 inhibition are still needed (and are underway). The high expression levels of AC1 in the hippocampus suggests that the initial focus of these studies should be on learning and memory. ST034307 is selective for AC1 vs. AC8. Nevertheless, AC1-KO mice still display impaired performance in certain learning and memory tasks.^4,16,17^ The use of a pharmacological agent will allow us to determine if those effects are a result of developmental issues (as AC1 expression is important for synaptic plasticity and development)^45,46^ or if there is an acute dose-dependent effect. If ST034307 is to be used for those experiments, intrathecal or intracerebroventricular injections will be required to ensure that the compound reaches the brain. The development of chronic adverse effects, other than analgesic tolerance, should also be investigated. It is not expected that AC1 inhibitors will be rewarding, but the current state of the opioid crisis indicates that this should be tested experimentally, and the effects of AC1 inhibitors on the release of dopamine in the nucleus accumbens should also be assessed.

It is noteworthy that the current experiments were performed with subcutaneous injections, instead of the intrathecal injections from Brust et al., 2017.^4^ This allowed us to determine the disposition of the compound in plasma and brain. The plasma concentration of 10 mg/kg ST034307 peaked at 60 minutes after injection. Notably, we were unable to detect ST034307 in the brain. Nevertheless, the disposition of this compound in plasma indicates a wide distribution in the body and rapid clearance resulting in relatively low concentrations compared to the administered dose. These concentrations persist for at least four hours to account for the effects that are seen in these experiments. These data suggest that the effects observed in the present studies are due to the actions of the compound in the periphery. It has recently been reported that the analgesic effects related to a reduction of AC1 activity is associated with the expression of the enzyme in the DRG and spinal cord.^9^ As the DRG is located in the periphery, it is likely that this is the site responsible for the analgesic properties of ST034307. The fact that ST034307 is not reaching the brain also precludes the compound’s activity in the hippocampus and makes it unlikely that this particular compound, when administered subcutaneously, would cause adverse effects related to learning and memory.

In the last set of data presented in the manuscript, the interaction between ST034307 and AC1 was mapped using molecular docking. Those results, achieved using of two different programs, suggest that ST034307 interacts at a site located between the ATP and forskolin binding sites. This binding site is located at the interface of the C1a and C2a domains and is indicative of a non-competitive or uncompetitive mechanism. The action of ST034307 is proposed to cause a disruption of the structure of AC1’s catalytic domain and, consequently, enzymatic inhibition. As our modeling showed that ST034307 does not bind to the ATP binding site, it is consistent with our previous findings that indicate that the compound is not a P-site inhibitor.^1,4^

As encouraging as the data presented in the manuscript appears, other compounds that looked promising in pre-clinical models of pain have failed to translate to clinic.^36^ While the nesting experiments account for some adverse effects and competing behaviors that may generate false positives, additional studies on ST034307 and the class of AC1 inhibitors are still needed. Particular attention should be devoted to possible impairments on learning and memory as well as other models of pain that reflect pain states that are different from the ones already examined. Experiments with AC1 inhibitors that can reach the brain are also desired. Nevertheless, the present work clearly demonstrates a correlation between selective inhibition of AC1 and behaviors that are consistent with analgesia in mice. More work is still needed to establish this class of compounds as novel pain therapeutics; however, the present study represents an important step that may signal that selective AC1 inhibitors should be prioritized for further testing and advancement for the treatment of pain.

## Supplementary Materials

Fig. S1. Ramachandran plot statistics analysis of the adenylyl cyclase 1 model.

Fig. S2. Main-chain stereochemical parameters statistical analysis of the adenylyl cyclase 1 model.

Fig. S3. Side-chain stereochemical parameters statistical analysis of the adenylyl cyclase 1 model.

Fig. S4. Three-dimensional profile analysis of the adenylyl cyclase 1 model.

Fig. S5. QMEANDisCo analysis of the adenylyl cyclase 1 model.

Fig. S6. 2D representation of the ST034307 poses.

## Data and materials availability

All data required to evaluate the conclusions in the paper are present in the manuscript or references. Materials will be made available upon request.

## Funding

This work is supported by the American Association of Colleges of Pharmacy’s New Investigator Award (T.F.B.), the Lloyd L. Gregory School of Pharmacy’s IntegraConnect grant (T.F.B and A.N.), and Palm Beach Atlantic University’s Quality Initiative grant (T.F.B.), Fundação Carlos Chagas Filho de Amparo à Pesquisa do Estado do Rio de Janeiro (FAPERJ) (N.B.), Conselho Nacional de Desenvolvimento Científico e Tecnológico (CNPq), Fundação de Apoio à Fiocruz (FIOTEC) (N.B.), Coordenação de Aperfeiçoamento de Pessoal de Nível Superior (CAPES) – Finance Code 001 (N.B.), and Programa Nacional de Apoio ao Desenvolvimento da Metrologia, Qualidade e Tecnologia (PRONAMETRO) from Instituto Nacional de Metrologia, Qualidade e Tecnologia (INMETRO) (T.V.A.F.).

## Author Contributions

Conceptualization: T.F.B. Experimental design: T.F.B., G.G., A.N., L.V.B.H. Performed experiments: T.F.B., G.G., T.P., L.V.B.H., A.S.M., T.V.A.F., V.S.P. Data analyses: T.F.B., L.V.B.H., A.N. Provided needed equipment or materials: N.B., H.W. Writing: T.F.B. wrote the first draft of the manuscript and all authors contributed, reviewed, and edited.

## Competing Financial Interests statement

The authors have no competing interests.

## Notes

### Competing Interest Statement

The authors have declared no competing interest.

