## Supplemental materials for "A selective adenylyl cyclase 1 inhibitor relieves pain without causing tolerance"

**This PDF file includes:**

Fig. S1. Ramachandran plot statistics analysis of the adenylyl cyclase 1 model.
Fig. S2. Main-chain stereochemical parameters statistical analysis of the adenylyl cyclase 1 model.
Fig. S3. Side-chain stereochemical parameters statistical analysis of the adenylyl cyclase 1 model.
Fig. S4. Three-dimensional profile analysis of the adenylyl cyclase 1 model.
Fig. S5. QMEANDisCo analysis of the adenylyl cyclase 1 model.
Fig. S6. 2D representation of the ST034307 poses.

#### Plot statistics

| | | |
|---|---|---|
| Residues in most favoured regions [A,B,L] | 302 | 88.0% |
| Residues in additional allowed regions [a,b,l,p] | 29 | 8.5% |
| Residues in generously allowed regions [~a,~b,~l,~p] | 7 | 2.0% |
| Residues in disallowed regions | 5 | 1.5% |
|  | ---- | ------ |
| Number of non-glycine and non-proline residues | 343 | 100.0% |
| Number of end-residues (excl. Gly and Pro) | 7 | |
| Number of glycine residues (shown as triangles) | 32 | |
| Number of proline residues | 9 | |
|  | ---- |  |
| Total number of residues | 391 | |

Based on an analysis of 118 structures of resolution of at least 2.0 Angstroms and R-factor no greater than 20%, a good quality model would be expected to have over 90% in the most favoured regions.

**Figure. S1.**

Ramachandran plot statistics analysis of the adenylate cyclase 1 (AC1) model made on the SAVES server using PROCHECK.

PROCHECK

### Main-chain parameters

#### saves

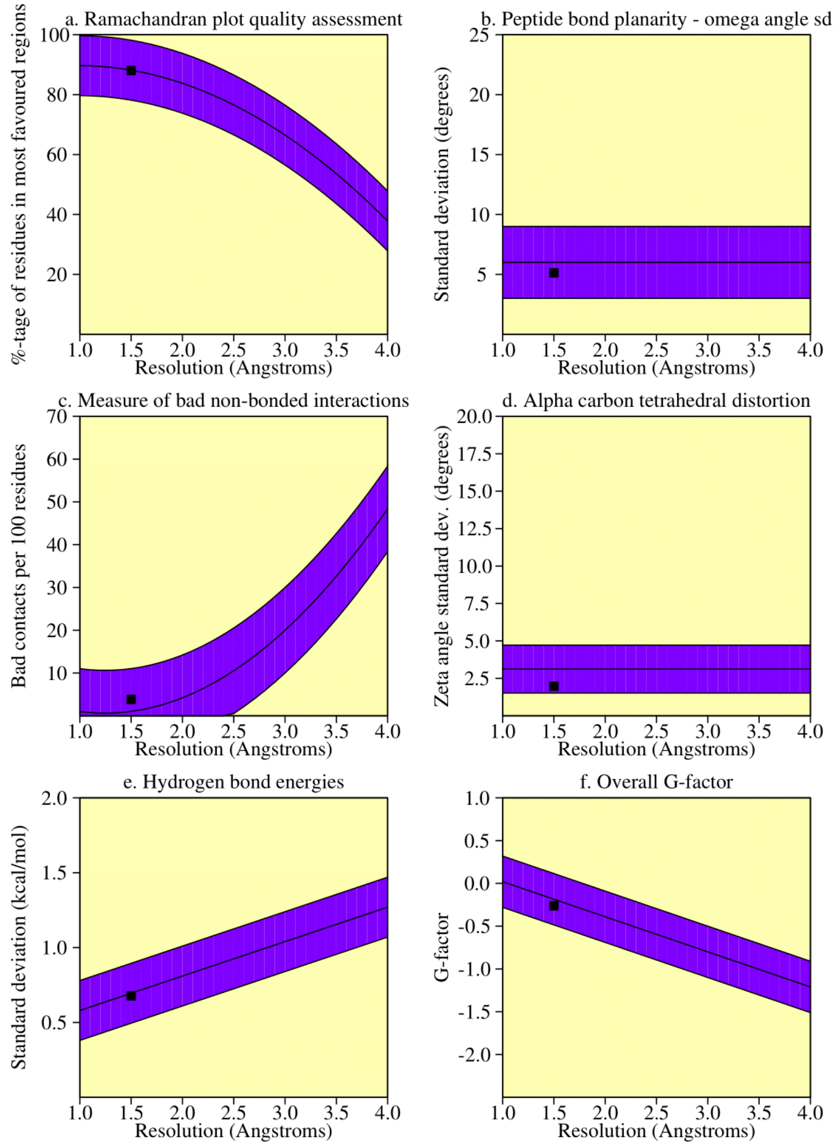


##### Plot statistics

| Stereochemical parameter | No. of data pts | Parameter value | Comparison values Typical value | Band width | No. of band widths from mean | |
|---|---|---|---|---|---|---|
| a. %-tage residues in A, B, L | 343 | 88.0 | 88.2 | 10.0 | -0.0 | Inside |
| b. Omega angle st dev | 385 | 5.1 | 6.0 | 3.0 | -0.3 | Inside |
| c. Bad contacts / 100 residues | 15 | 3.8 | 1.0 | 10.0 | 0.3 | Inside |
| d. Zeta angle st dev | 355 | 2.0 | 3.1 | 1.6 | -0.7 | Inside |
| e. H-bond energy st dev | 256 | 0.7 | 0.7 | 0.2 | -0.1 | Inside |
| f. Overall G-factor | 391 | -0.3 | -0.2 | 0.3 | -0.3 | Inside |

**Figure. S2.**

Main-chain stereochemical parameters statistical analysis of the AC1 model made on the SAVES server using PROCHECK.

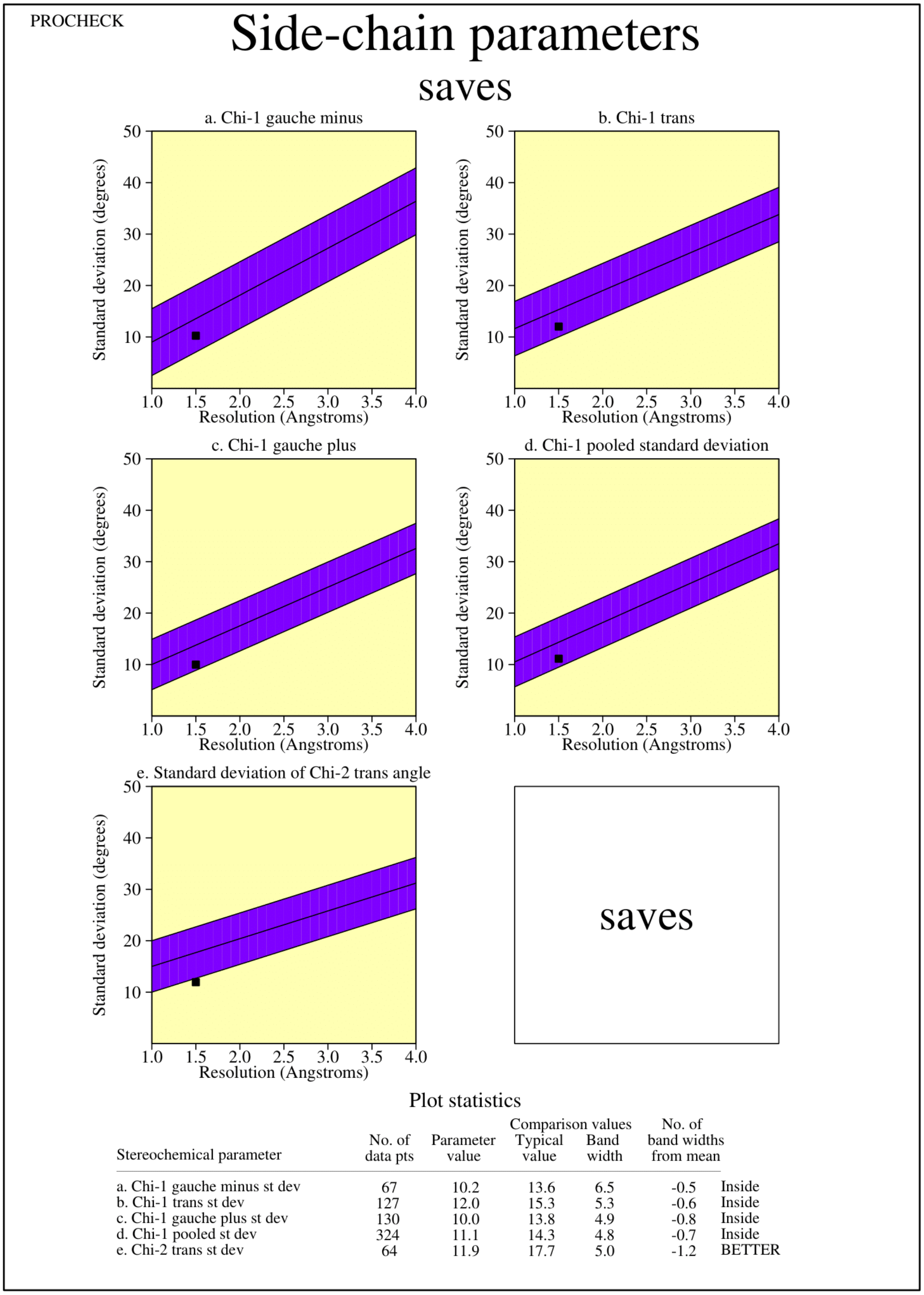


| Stereochemical parameter | No. of data pts | Parameter value | Comparison values Typical value | Band width | No. of band widths from mean | |
|---|---|---|---|---|---|---|
| a. Chi-1 gauche minus st dev | 67 | 10.2 | 13.6 | 6.5 | -0.5 | Inside |
| b. Chi-1 trans st dev | 127 | 12.0 | 15.3 | 5.3 | -0.6 | Inside |
| c. Chi-1 gauche plus st dev | 130 | 10.0 | 13.8 | 4.9 | -0.8 | Inside |
| d. Chi-1 pooled st dev | 324 | 11.1 | 14.3 | 4.8 | -0.7 | Inside |
| e. Chi-2 trans st dev | 64 | 11.9 | 17.7 | 5.0 | -1.2 | BETTER |

**Figure. S3.**

Side-chain stereochemical parameters statistical analysis of the AC1 model made on the SAVES server using PROCHECK.

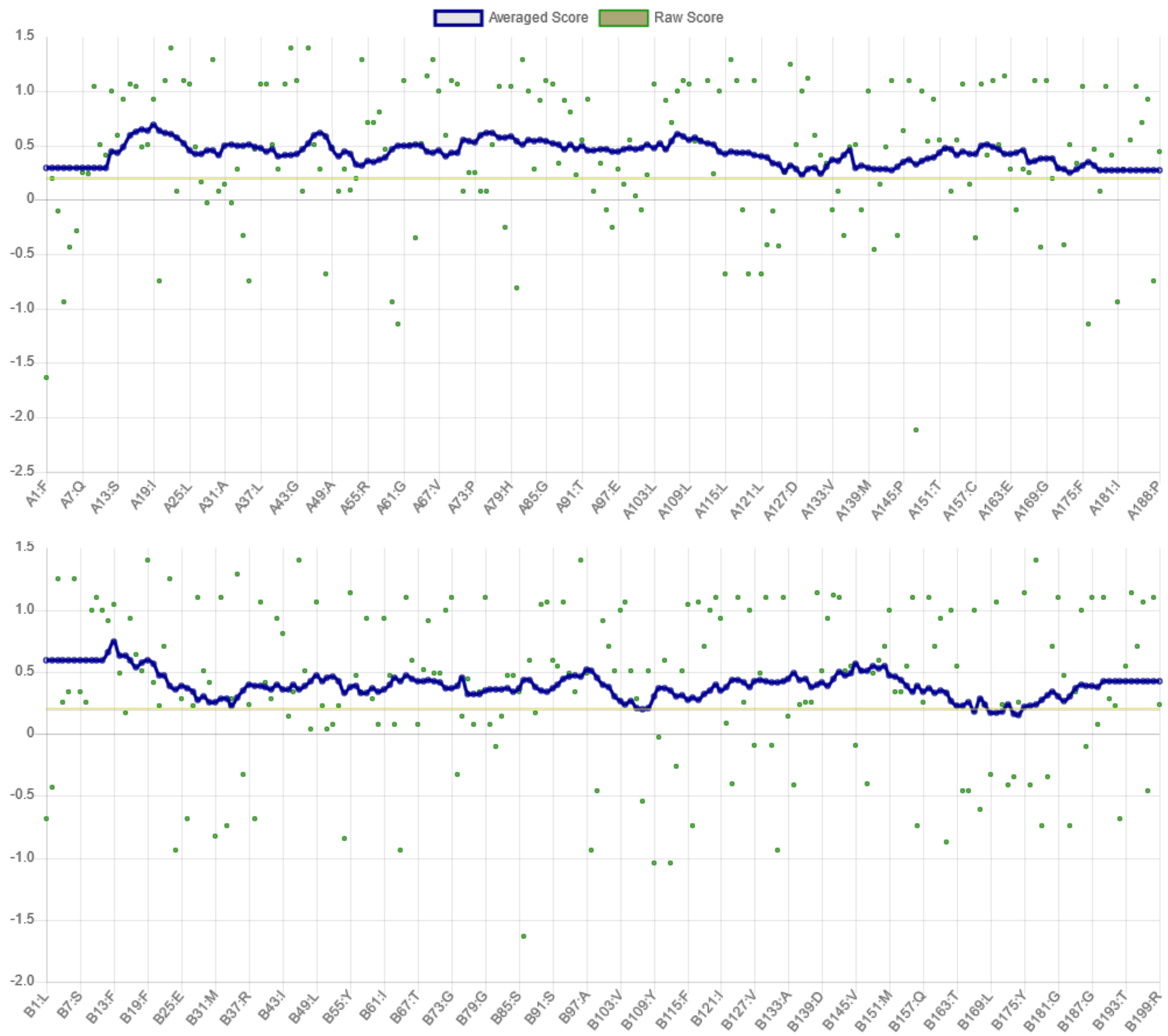


**Figure S4**

Three-dimensional profiles analysis of the AC1 model made on the SAVES server using VERIFY 3D for chain C1a and C2a, respectively. 98.45% of the residues have averaged 3D-1D score >= 0.2**.**

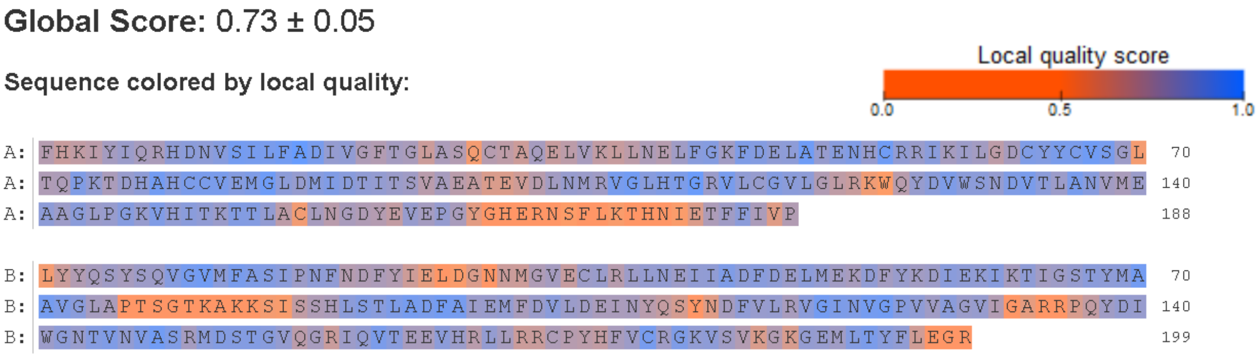


**Figure S5**

QMEANDisCo analysis of the AC1 model made on the SWISS-PROT server. QMEANDisCo scored regions of the model from 0 to 1, or poor to good, based on statistical potentials of mean force and agreement terms with consensus-based distance constraints.

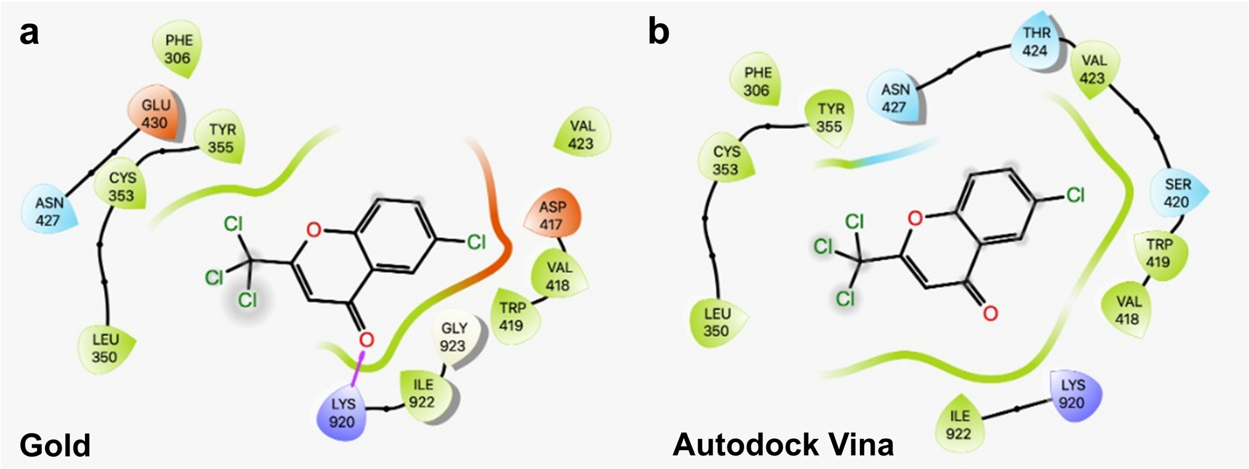


**Figure S6**

2D representation of the ST034307 poses, using Gold (A) and Autodock Vina (B) programs, showing the hydrogen bond (purple arrows) and steric interactions with the AC1 model.
